# IBA endogenous auxin regulates Arabidopsis root system development in a glutathione-dependent way and is important for adaptation to phosphate deprivation

**DOI:** 10.1101/818344

**Authors:** José A. Trujillo-Hernandez, Laetitia Bariat, Lucia C. Strader, Jean-Philippe Reichheld, Christophe Belin

## Abstract

Root system architecture results from a highly plastic developmental process to perfectly adapt to environmental conditions. In particular, the development of lateral roots (LR) and root hair (RH) growth are constantly optimized to the rhizosphere properties, including biotic and abiotic constraints. Every step of root system development is tightly controlled by auxin, the driving morphogenic hormone in plants. Glutathione, a major thiol redox regulator, is also critical for root system development but its interplay with auxin is still scarcely understood. Indeed, previous works showed that glutathione deficiency does not alter root responses to exogenous indole acetic acid (IAA), the main active auxin in plants. Because indole butyric acid (IBA), another endogenous auxinic compound, is an important source of IAA for the control of root development, we investigated the crosstalk between glutathione and IBA during root development. We show that glutathione deficiency alters LR and RH responses to exogenous IBA but not IAA. Although many efforts have been deployed, we could not identify the precise mechanism responsible for this control. However, we could show that both glutathione and IBA are required for the proper responses of RH to phosphate deprivation, suggesting an important role for this glutathione-dependent regulation of auxin pathway in plant developmental adaptation to its environment.

## INTRODUCTION

The plasticity of root system development, combining root growth and root branching, is essential for plants to sustain, adapt and optimize their growth in changing environmental conditions, such as nutrient and water availability, rhizosphere microbiome or soil structure heterogeneity. Root growth relies on the activity of the root apical meristem that regulates histogenesis via the control of cell proliferation. Root branching is a complex organogenesis process that allows the development of new lateral roots (LR) from regularly spaced pericycle founder cells in *Arabidopsis thaliana*. These founder cells are originally specified by an oscillatory mechanism occurring in the root tip and involving cyclic programmed cell death of lateral root cap cells (Moreno-Risueno et al., 2010; Xuan et al., 2016). In addition to the root system architecture, the development of epidermal root hairs (RH) is particularly sensitive to changes in environmental conditions and contribute to the root system adaptation (Grierson et al., 2014).

Among nutrients, phosphorus has a key role in all living organisms and is one of the main limiting factors for plant growth and crop productivity (Leinweber et al., 2018). Its homeostasis is highly dependent on phosphate uptake by the root cap (Kanno et al., 2016). Phosphate concentration strongly impacts root system development in many plant species including Arabidopsis (López-Bucio et al., 2002, 2003; Hodge, 2004). In particular, general low phosphate availability is well known to increase RH length (Bates and Lynch, 1996). Biotic factors also modulate root development. Among them, rhizospheric PGPR (plant growth promoting rhizobacteria) are able to stimulate RH elongation (Poitout et al., 2017).

Auxins, and particularly the most abundant endogenous one, indole acetic acid (IAA), play a pivotal and integrative role in all steps of root system development, both under optimal or stress conditions (Saini et al., 2013; Korver et al., 2018). During LR development, auxin is responsible for the activation of founder cells, the development and organization of the LR primordia, and the emergence of the newly formed LR through the external layers of the primary root. Even earlier, the oscillatory positioning of pericycle founder cells depends on auxin maxima generated via auxin release by dying root cap cells (Moreno-Risueno et al., 2010; Du and Scheres, 2018). Auxin also modulates RH elongation, which has been shown to mediate responses to stimuli such as abscisic acid, PGPR or phosphate deprivation (Wang et al., 2017; Poitout et al., 2017; Nacry et al., 2005). Recently, a set of works on Arabidopsis and rice evidenced that RH response to phosphate deprivation requires an increase in auxin *de novo* synthesis in the root cap, and its apico-basal transport to the epidermal cells via the auxin influx facilitator AUX1 (Giri et al., 2018; Bhosale et al., 2018; Parry, 2018).

Indole-3-butyric Acid (IBA) is a structural derivative of IAA, differing by only two additional carbons in the side chain (Korasick et al., 2013). Although convincing evidence support a role for IAA in IBA biosynthesis, the mechanisms responsible and enzymes involved are still unknown (Ludwig-Müller et al., 1995; Frick and Strader, 2018). It is now broadly accepted that IBA solely acts as an IAA precursor through its β-oxidative decarboxylation in peroxisomes (reviewed in Frick and Strader, 2018). This enzymatic process involves several enzymes, some shared with other β-oxidation pathways such as PED1, others apparently specific to the IBA-to-IAA conversion, such as IBR1, IBR3, IBR10 and ECH2 (Strader and Bartel, 2011; Frick and Strader, 2018). IBA homeostasis is also regulated via its transport and conjugation, but only few regulators have been identified. Type G ABC transporters ABCG36 and ABCG37 can efflux IBA from the cells (Strader and Bartel, 2009; Ruzicka et al., 2010), the NPF family member TRANSPORTER OF IBA1 transports IBA across the vacuolar membrane (Michniewicz et al., 2019), while the generic PXA1/COMATOSE transporter is likely to import IBA into the peroxisome. Like other auxins, IBA is known to conjugate with sugars and amino acids. UGT84B1, UGT74D1, UGT74E2 and UGT75D1 can conjugate IBA to glucose (Jackson et al., 2001; Tognetti et al., 2010; Jin et al., 2013; Zhang et al., 2016), whereas GH3.15 has recently been shown to be able to conjugate IBA to amino acids (Sherp et al., 2018). IBA-derived IAA plays important roles during root development, including root apical meristem maintenance and adventitious rooting. IBA conversion to IAA is also critical for RH elongation (Strader et al., 2010). Finally, recent works have reported the critical role of IBA-to-IAA conversion in the root cap as a source of auxin for the oscillatory positioning of LR founder cells (De Rybel et al., 2012; Xuan et al., 2015). IBA-to-IAA conversion taking place in the LR primordium itself also likely participates in further LR development (Strader and Bartel, 2011; Michniewicz et al., 2019). Despite all these reported functions in root development, the importance of IBA-derived IAA relatively to other IAA sources is still poorly understood and scarcely documented, particularly in case of changing environmental constraints.

Changes in environmental conditions result in Reactive Oxygen Species (ROS) imbalance, thus affecting the general redox cellular homeostasis. For example, phosphate deficiency alters H_2_O_2_ and O_2_^・-^ production in roots (Tyburski et al., 2009). ROS modulate many aspects of plant development, including root system development (Singh et al., 2016; Tsukagoshi, 2016). Controlled ROS production by NADPH oxidases are particularly involved in RH elongation and LR development (Orman-ligeza et al., 2016; Mangano et al., 2016; Carol and Dolan, 2006; Mangano et al., 2018). Auxin and ROS pathways interact to control root development, since auxin can trigger ROS production and in turn ROS can affect IAA levels and transport (Orman-ligeza et al., 2016; Tognetti, 2017; Zwiewka et al., 2019).

Although ROS are important signalling molecules, they are also potent oxidants that can damage proteins, lipids or nucleic acids. Organisms have evolved many antioxidant systems to prevent or repair such damages (Noctor et al., 2018). Among them are several detoxifying enzymes such as catalases or peroxidases, able to reduce the most stable ROS, hydrogen peroxide (H_2_O_2_), but also several antioxidant molecules such as ascorbate or glutathione. Glutathione is a small and stable thiol-containing tripeptide (Glu-Cys-Gly) essential for plant survival and present in large concentrations in cells, up to several millimolar (Noctor et al., 2012). It has many roles in almost all cellular compartments, including the detoxification of heavy metals and xenobiotics, sulfur homeostasis, ROS homeostasis and redox signalling. To ensure these functions, glutathione can act as a precursor (e.g. phytochelatins), be conjugated to other molecules via the Glutathione-S-transferase enzyme family, or serve as an electron donor for antioxidant systems (Noctor et al., 2012). Reduced glutathione (GSH) is therefore converted to oxidised glutathione (GSSG) which is reduced back by glutathione reductases. In standard growth conditions GSH is in large excess relative to GSSG. Glutathione biosynthesis is a 2-step reaction, with *GSH1* encoding the rate-limiting γ-glutamylcysteine synthetase and *GSH2* encoding a glutathione synthetase (Noctor et al., 2012). Knock out mutations in any of these two genes are lethal. However, genetic screens identified several knock-down alleles of *GSH1*, allowing to get plants with reduced glutathione levels (Vernoux et al., 2000; Ball et al., 2004; Jobe et al., 2012; Howden et al., 1995; Parisy et al., 2007; Shanmugam et al., 2012). The importance of glutathione in root development is illustrated by the absence of root meristem maintenance in *rml1* mutant, together with the impairment of primary root growth and LR development in less severe *gsh1* alleles (Bashandy et al., 2010; Koprivova et al., 2010; Marquez-Garcia et al., 2014; Vernoux et al., 2000).

We previously reported that *cad2* or *pad2* mutants can respond almost normally to exogenous IAA for root development (Bashandy et al., 2010), although auxin transport seems to be affected in glutathione-deficient plants (Koprivova et al., 2010; Bashandy et al., 2010). The role of glutathione in controlling root development is therefore still misunderstood. Given the importance of IBA-derived IAA in regulating root development, we chose to address the role of glutathione in root responses to IBA. We show that glutathione deficiency impairs LR and RH responses to IBA but not IAA. We then try to identify the glutathione-dependent mechanisms required for IBA responses but every IBA-related pathway examined does not reveal sensitivity to glutathione levels. We finally suggest a physiological role for this glutathione-dependent IBA response, showing that IBA and glutathione pathways are critical for RH responses to phosphate deficiency.

## RESULTS

### Glutathione deficiency specifically alters LR and RH responses to IBA but not IAA

Since IBA plays important roles during root development, we investigated root responses to both IAA and IBA in several *gsh1* weak alleles and in plants treated with Buthionine Sulfoximine (BSO), a chemical specific inhibitor of GSH1 enzyme.

We first wanted to precisely know the glutathione levels in the different genotypes in our growth conditions, thus we quantified endogenous glutathione in whole 8-day old seedlings (Figure 1A). We find the same amount of total glutathione (*i.e.* about 25% of wild-type content) in *cad2*, *pad2* and *zir1* mutants, which was expected for *cad2* and *pad2* whereas *zir1* has previously been reported to have lower glutathione contents (about 15% relative to wild-type). We show that 1mM exogenous BSO also reduces endogenous glutathione levels of wild type plants to approximately the same levels. In addition, we report that exogenous IBA treatment does not impact endogenous glutathione levels.

**Figure 1.**
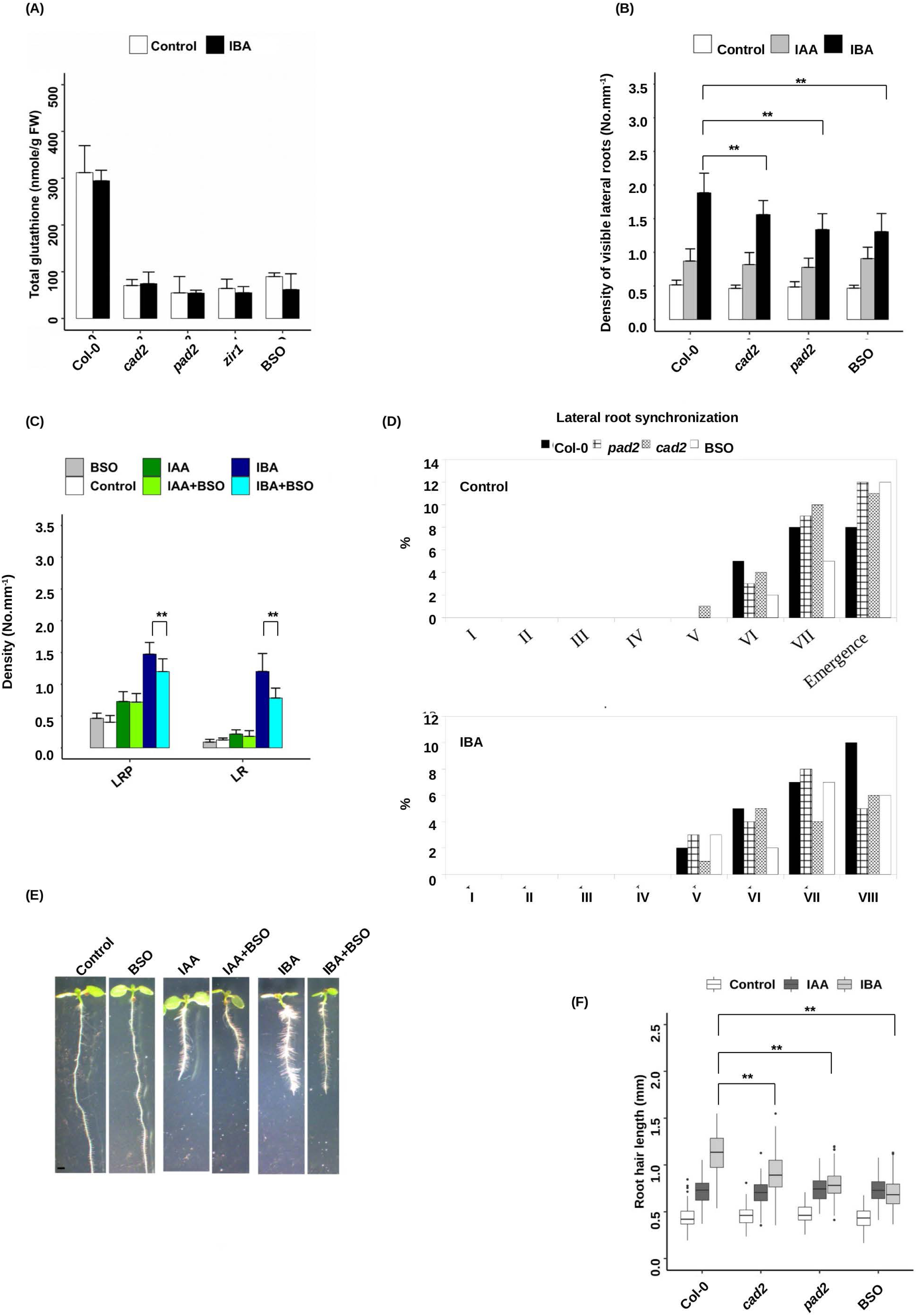
Glutathione specifically regulates root responses to IBA. A. Total glutathione content in 8-day-old wild-type (Col-0), *cad2*, *pad2* or *zir1* seedlings grown on standard ½ MS medium. Wild-type plants grown in the presence of 0.5 mM BSO were also assayed (BSO). Data represent the means of 3 biological repetitions. B. Emerged lateral root density of 10-day-old wild-type (Col-0), *cad2* or *pad2* plants grown on standard ½ MS medium and wild-type plants grown in the presence of 0.5 mM BSO (n>15). C. Emerged LR and LR primordia density of 10-day-old wild-type plants grown on standard ½ MS medium (control), or in the presence of different combinations of BSO 0.5 mM, IBA 10 µM and/or IAA 50 nM, as mentioned (n>15). D. Developmental stages of LR primordia of 6-day-old wild-type (Col-0), *cad2* or *pad2* seedlings 48 hours after gravistimulation on standard ½ MS medium; wild-type plants grown in the presence of 0.5 mM BSO were also assayed (BSO). Data indicate the percentage of LRP in each developmental stage (n> 16) and are representative of 3 independent experiments. E. Pictures of representative 8-day-old wild-type plants grown on standard ½ MS medium, or in the presence of different combinations of BSO 0.5 mM, IBA 10 µM and/or IAA 50 nM, as mentioned. F. Quantification of root hair length of 8-day-old wild-type (Col-0), *cad2* or *pad2* mutant plants grown on standard ½ MS medium (Control), or in the presence of 50 nM IAA (IAA) or 10 µM IBA (IBA). Wild-type plants in the presence of 0.5 mM BSO (BSO) were also assayed (n>100). Histograms represent the mean and error bars represent the standard deviation. Asterisks indicate a significant difference, based on a two-tailed Student t-test (** P≤0.001).

Supplemental Figures 1A and 1B show that *cad2* and *pad2* mutants display the same primary root growth as the wild-type in our growth conditions, and that the primary root responds normally to both IAA and IBA. In the same way, BSO treatment (1mM) neither affects primary root growth nor its response to IAA or IBA. *zir1* mutant shows a strongly reduced primary root growth under normal conditions but responds normally to both IAA and IBA. Since we measured the same total glutathione levels in *zir1* than in *cad2* or *pad2* but observe a severely reduced growth of that allele, we decided to focus on *pad2*, *cad2* and BSO treatment in the next experiments.

As expected, both IAA and IBA also induce LR density in the mature zone of the root in 10-day-old seedlings (Figure 1B). LR density in both *cad2* and *pad2* mutants responds normally to IAA treatment but displays hyposensitivity to exogenous IBA. Interestingly, the addition of 1 mM BSO phenocopies *cad2* and *pad2* mutants, and this BSO phenotype is dose-sensitive (Supplemental Figure 2A). Finally, we could revert *cad2* hyposensitivity to IBA by adding exogenous GSH (Supplemental Figure 2B). Because emerged LR density depends both on LR initiation and subsequent LRP development, we also addressed the LRP density in the same conditions. Figure 1C reveals that both LRP and emerged LR densities display IBA hyposensitivity in plants with low glutathione levels, suggesting that glutathione affects LR development upstream of LRP development. Another way to investigate LRP development is to force and synchronize LRP initiation by gravistimulation (Péret et al., 2012). We therefore gravistimulated glutathione-deficient plants, in the absence or presence of exogenous IBA (1 mM). We first observe that exogenous IBA treatment slightly slows down wild-type LRP development upon gravistimulation in our growth conditions (Figure 1D). We also observe that *cad2*, *pad2* and BSO-treated plants behave exactly like wild-type in such an assay. Taken together, these results tend to support a role for glutathione in IBA-dependent specification or activation of founder cells prior to the LR development process.

Finally, we monitored RH elongation responses to IBA and IAA in glutathione-deficient plants (Figure 1E and 1F). As for LR, we notice that *cad2* and *pad2* mutants are hyposensitive to the IBA-dependent induction of RH growth but respond normally to IAA. Similarly, 1 mM BSO treatment also reduces the RH response to IBA but not IAA. We can therefore conclude that glutathione is also required for the IBA-dependent induction of RH growth.

### Glutathione levels affect auxin signalling in the basal part of the meristem

We have shown that glutathione deficiency alters RH elongation and founder cells specification or activation in response to IBA. We know that root tip-derived IAA transits through the lateral root cap to regulate both founder cell positioning and RH growth in the basal part of the meristem. We therefore investigated auxin response in the root tip, by using the *DR5:GUS* auxin signalling reporter (Ulmasov et al., 1997). In standard growth conditions, we only reveal GUS staining in the quiescent center and the columella, and BSO treatment by itself does not change the *DR5:GUS* expression pattern (Figure 2A). As expected, exogenous IAA treatment for 24 hours induces auxin response in the whole root tip and root epidermis, and the presence of BSO has again no effect on this distribution. In response to IBA, GUS staining increases in the distal meristem and strongly appears specifically in the trichoblast epidermal cell files in the basal part of the meristem, up to the differentiation zones. In the presence of BSO, this IBA-dependent strong signal in the basal meristem almost disappears while the signal in the distal meristem remains strong.

**Figure2.**
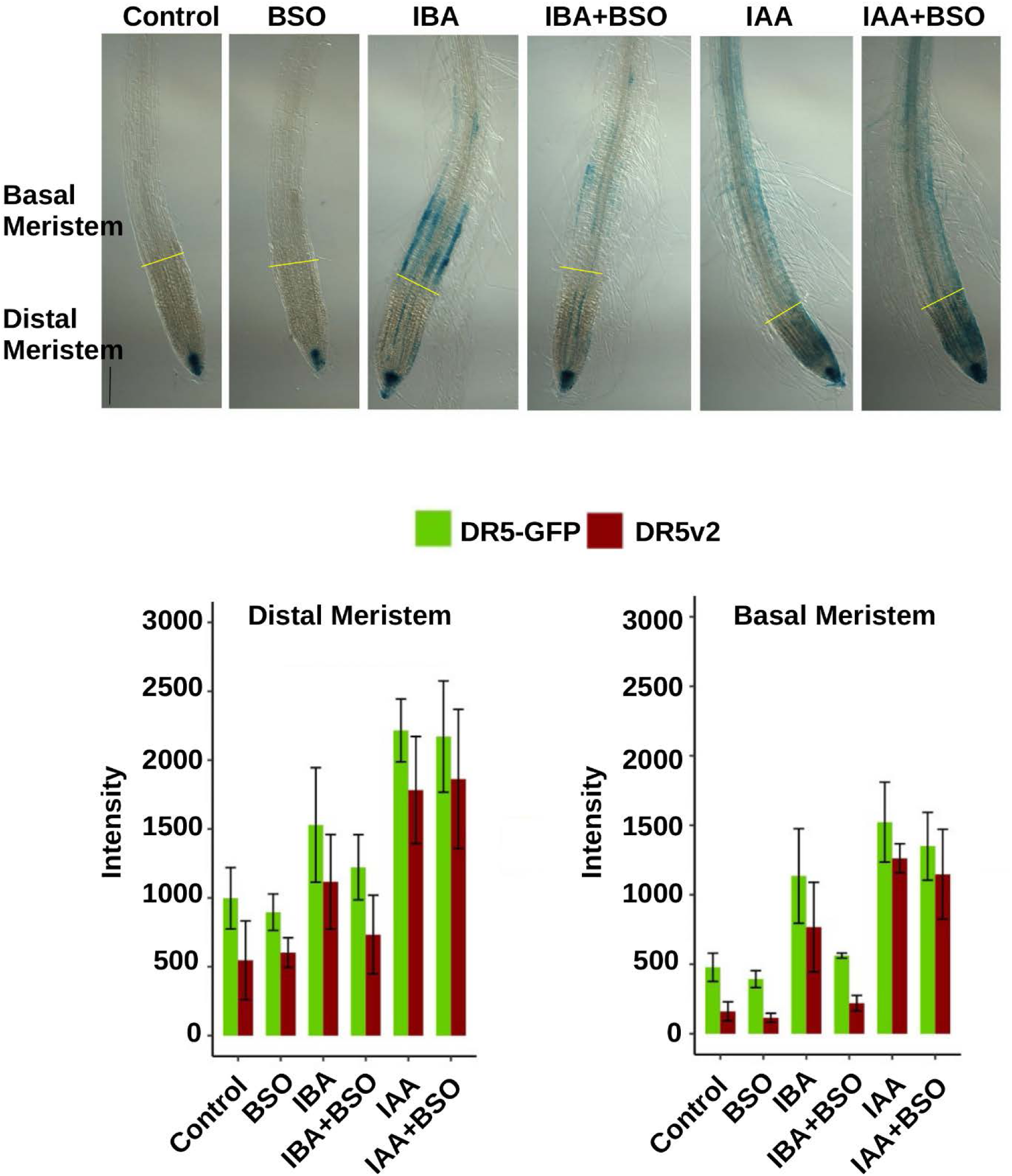
Glutathione is critical for the IBA-derived IAA signalling in the basal meristem. A. Representative Pictures of 8-day-old GUS-stained *DR5:GUS* plants grown on standard ½ MS medium (Control) or in the presence of 0.5 mM BSO (BSO), after a 24-hour long treatment with 10 µM IBA or 50 nM IAA as precised (n≥7). B. Quantification of GFP and Tomato fluorescent signals from 8-day-old expressing the *DR5:n3EGFP/DR5v2:ntdTomato* double reporter, and grown in the same conditions as A (n≥7).

In order to confirm these results, we used another auxin signalling marker that allows the quantification of auxin response, the *DR5:n3EGFP/DR5v2:ntdTomato* double reporter line (Liao et al., 2015). We quantified both GFP and Tomato fluorescence in the distal and basal parts of the meristem (Figure 2B). In agreement with the *DR5:GUS* reporter line, both IBA and IAA treatments increase auxin signalling in distal and basal parts of the meristem. The presence of BSO does not significantly alter auxin signalling, neither in standard conditions nor upon exogenous IAA treatment. However, we can confirm that BSO severely represses IAA signalling in the basal part of the meristem in response to exogenous IBA, while it has limited effect in the distal part.

Hence we show that glutathione is specifically required for IBA-derived IAA signalling in the basal part of the meristem, where LR founder cells are specified and RH growth is determined. Moreover, we show that glutathione does not affect IAA signalling components since auxin signalling reporters respond normally to exogenous IAA when glutathione content is depleted.

### IBA-derived IAA responses are specifically affected by glutathione deficiency

Because glutathione is a critical regulator of cellular redox homeostasis, we addressed root responses to IBA in other redox-related mutants. We chose to analyse the *gr1* mutant, affected in cytosolic and peroxisomal glutathione reduction, the *cat2* mutant, affected in H_2_O_2_ detoxification, and the *ntra ntrb* double mutant affected in the thioredoxin-dependent thiols reduction system. Quantification of both GSH and GSSG (Figure 3A) shows that *gr1*, *ntra ntrb* and *cat2* mutant plants have higher total glutathione concentrations than wild type plants. The application of exogenous IBA has almost no effect. As expected, *cat2* and *gr1* mutants also display higher glutathione oxidation status. In contrast to *cad2*, none of the other mutants display any LR density hyposensitivity to IBA (Figure 3B). In addition, the use of the roGFP2 probe allowed us to confirm that the presence of IBA does not generate any imbalance in glutathione redox status in root tissues (Figure 3C and Supplemental Figure 3). These results suggest that root responses to IBA specifically depend on glutathione overall levels rather than glutathione redox status or any general redox imbalance.

**Figure 3.**
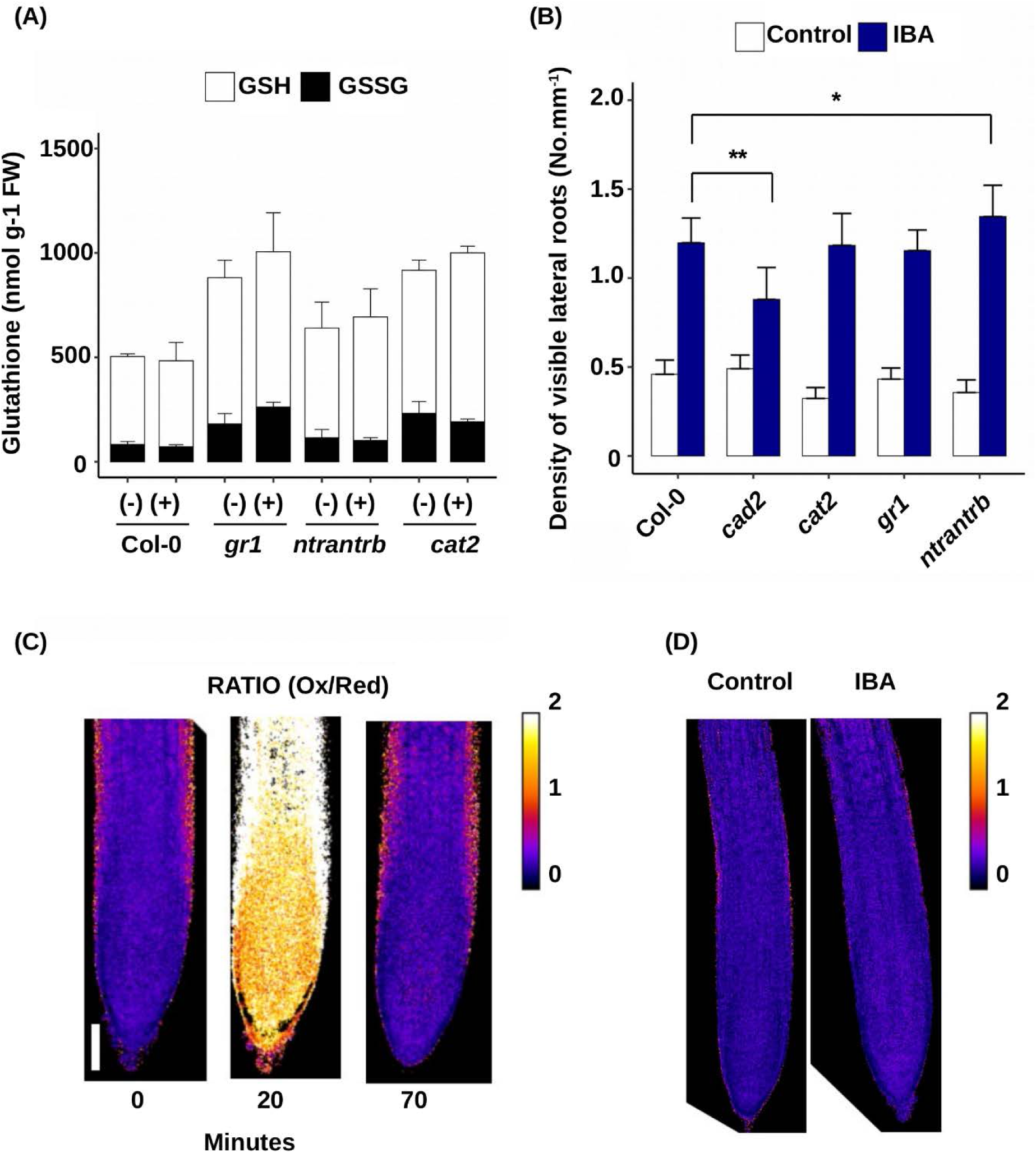
IBA hyposentitivity phenotype is specific to GSH-depleted plants. A. Glutathione content in 8-day-old wild-type (Col-0), *cad2*, *gr1*, *ntra ntrb* or *cat2* seedlings grown on standard ½ MS medium. Data represent the means of 3 biological repetitions. Reduced glutathione is represented in white (GSH) and oxidised glutathione in black (GSSG). B. Emerged lateral root density of 10-day-old wild-type (Col-0), *cad2*, *cat2*, *gr1* or *ntra ntrb* plants grown on standard ½ MS medium or the presence of 10 µM IBA (n>16). C. Representative pictures (n>10) of roGFP2 reporter line grown on standard ½ MS medium (MS), submitted to a 30 min treatment with 1 mM H_2_O_2_ then to an additional 30 min treatment with 10 mM DTT (top panel). The bottom panel represents roGFP2 reporter plants grown on standard ½ MS medium supplemented or not with 10 µM IBA. Pictures are made from the ratio between the oxidised roGFP2 signal (excitation at 405 nm) and the reduced roGFP2 signal (excitation at 488 nm). Ratio values are represented in the color scale. Histograms represent the mean and error bars represent the standard deviation. Asterisks indicate a significant difference, based on a two-tailed Student t-test (*P<0.01; **P<0,001).

### IAA transport from the distal to the basal meristem is not targeted by glutathione

We have just shown that IBA-derived IAA response in the basal meristem is dependent on glutathione levels. We also know that AUX1 is required for IAA to regulate RH growth in response to phosphate deprivation and that IBA-to-IAA conversion in the root cap is required for LR founder cells specification in the basal meristem (Bhosale et al., 2018; Xuan et al., 2015). AUX1 auxin influx facilitator and PIN2 auxin efflux carrier are the IAA transporters that ensure the apico-basal IAA flux in the root cap and the root tip epidermis. Finally, we previously published that strong BSO treatment induces long-term decrease in the expression of several genes encoding IAA transporters, including AUX1 and PIN members.

In order to determine if AUX1 and/or PIN2 are the targets of glutathione to modulate root responses to IBA, we examined LR density in *aux1* and *pin2* mutants. As shown in Figure 4A, *aux1* and *pin2* mutants still display hyposensitivity to IBA upon BSO treatment. In other words, the glutathione regulation still occurs in both *aux1* and *pin2* mutants, revealing that neither AUX1 nor PIN2 is regulated by glutathione to control root responses to IBA. Because AUX1 and PIN2 are members of multigene families, we wanted to ensure that other members of AUX/LAX or PIN families are not replacing AUX1 and PIN2 in the respective mutants. We therefore addressed IBA responses with or without BSO treatment in the presence of specific inhibitors of both families (Figure 4B). We observed that BSO still leads to LR hyposentitivity to IBA both in the presence of *N*-1-naphthylphthalamic acid (NPA), that inhibits PIN-dependent auxin efflux, and 1-naphthoxyacetic acid (NOA), a specific inhibitor of AUX/LAX influx facilitators. All together these data suggest that the glutathione-dependent control of root responses to IBA does not affect IAA transport from the root apex to the basal meristem.

**Figure 4.**
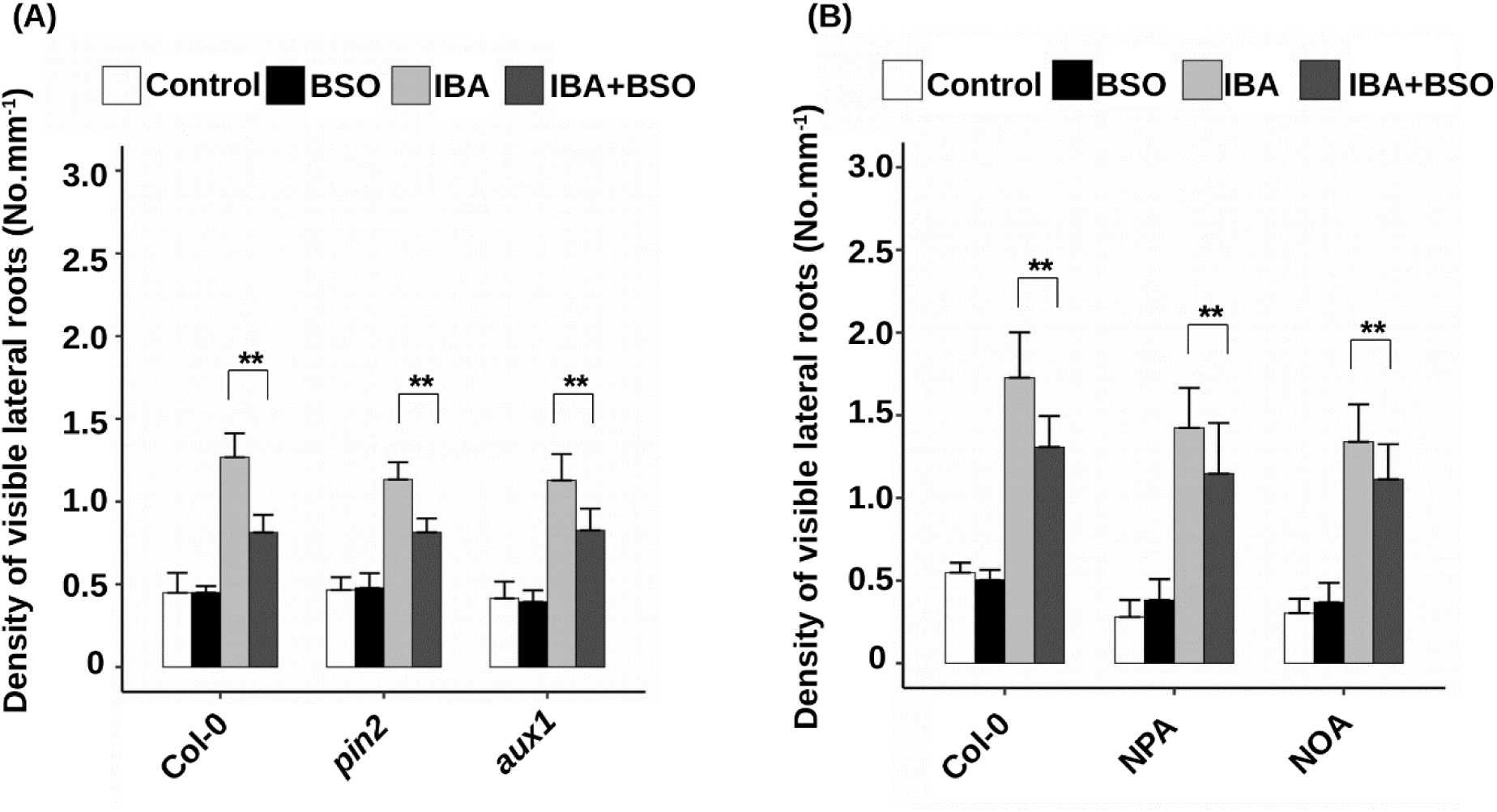
IAA transport is not the target of glutathione. A. Emerged lateral root density of 10-day-old wild-type (Col-0), *aux1* or *pin2* plants grown on standard ½ MS medium, or in the presence of different combinations of BSO 0.5 mM and IBA 10 µM (n>10). B. Emerged lateral root density of 10-day-old wild-type (Col-0) grown on standard ½ MS medium, supplemented or not with the transport inhibitors NPA 1 µM or NOA 1 µM (n≥15). Histograms represent the mean and error bars represent the standard deviation. Asterisks indicate a significant difference, based on a two-tailed Student t-test (**P<0,001).

### Looking for glutathione targets in IBA pathways

Since IAA transport was not the target of glutathione, we investigated IBA homeostasis in plants with low glutathione content.

Interestingly, *gsh1* mutants display pleiotropic phenotypes opposite to the phenotypes of mutant alleles in *PDR8/PEN3/ABCG36* for IBA sensitivity, but also for sensitivity to *Pseudomonas* and cadmium treatments (Kim et al., 2007; Lu et al., 2015; Strader and Bartel, 2009). ABCG36 is responsible for IBA efflux from the cells, which prompted us to investigate the IBA import into plant cells in the *cad2* mutant. As shown in Figure 5A, we did not detect any impairment in ^3^H-IBA accumulation in *cad2*. This suggests that IBA import into the cell is not perturbed by glutathione deficiency.

**Figure 5.**
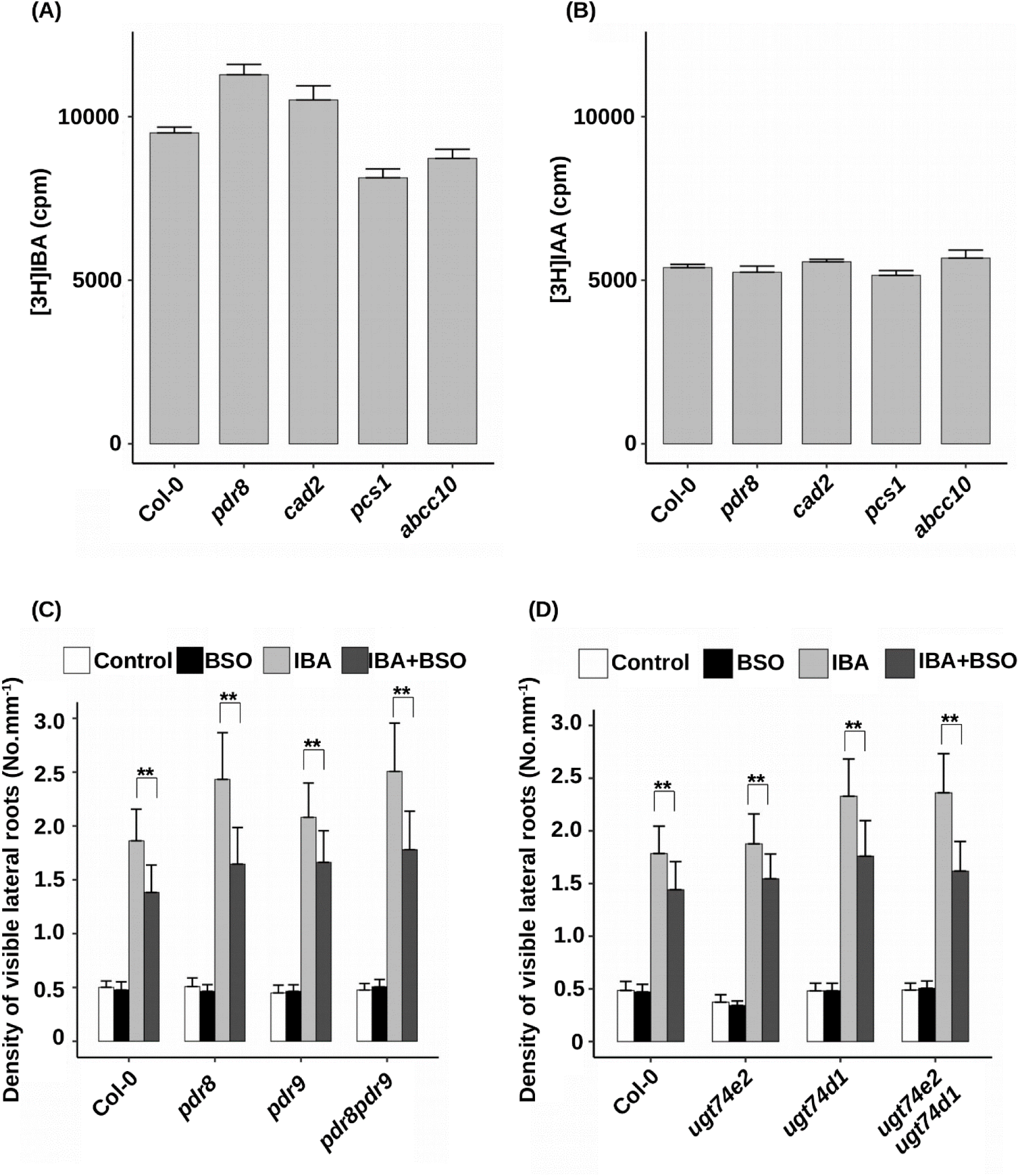
IBA homeostasis does not display sensitivity to glutathione. A. Uptake of [^3^H]IBA by wild type (Col), *pen3-4* (*pdr8*), *cad2-1*, *pcs1* or *abcc10* excised root tips of 8-day-old plants. Data represent eight replicates with five root tips per genotype and per replicate. B. Uptake of [^3^H]IAA by wild type (Col), *pen3-4* (*pdr8*), *cad2-1*, *pcs1* or *abcc10* excised root tips of 8-day-old plants. Data represent eight replicates with five root tips per genotype and per replicate. C. Emerged lateral root density of 10-day-old wild-type (Col-0), *pdr8*, *pdr9* or *pdr8 pdr9* plants grown on standard ½ MS medium, or in the presence of different combinations of BSO 0.5 mM and IBA 10 µM (n≥15). D. Emerged lateral root density of 10-day-old wild-type (Col-0), *ugt74e2*, *ugt75d1* or *ugt74e2 ugt75d1* plants grown on standard ½ MS medium, or in the presence of different combinations of BSO 0.5 mM and IBA 10 µM (n≥15). Histograms represent the mean and error bars represent the standard deviation. Asterisks indicate a significant difference, based on a two-tailed Student t-test (**P<0,001).

In order to better investigate IBA transport, we analysed the glutathione-dependent LR and RH response to IBA in *abcg36/pdr8* and *abcg37/pdr9* mutants (Figure 5B and Supplemental Figure 4). As expected, *pdr8* mutant displays hypersensitivity to exogenous IBA. However, both mutants are still clearly resistant to IBA in the presence of BSO. Because of putative redundancy between these two proteins, we generated the *pdr8 pdr9* double mutant that displays a *pdr8*-like hypersensitivity of LR density to exogenous IBA. As for single mutants, LR density and RH length in the double mutant are induced by IBA but BSO is still able to decrease this response. This result means that IBA efflux transporters ABCG36 and ABCG37 are not the targets of glutathione-dependent regulation.

We know that IBA homeostasis is also regulated via conjugation with glucose, and several glycosyltransferases are able to catalyse such a reaction (Jackson et al., 2001; Tognetti et al., 2010; Jin et al., 2013; Zhang et al., 2016). Among them, we assayed *ugt74d1* and *ugt74e2* mutants. Again, we could observe that LR and RH responses to IBA are still BSO-sensitive in both mutants, although *ugt74d1* LR density is hypersensitive to IBA compared to wild-type (Figure 5C and Supplemental Figure 4). Because of putative redundancy between these two glycosyltransferases, we generated an *ugt74e2 ugt74d1* double mutant. However, the response to IBA is still reduced in the presence of BSO in the double mutant, suggesting that these glycosyltransferases are not the targets of glutathione-dependent regulation.

Finally, we also investigated the enzymatic pathway involved in the IBA-to-IAA conversion in the peroxisome. Figure 6A shows that *ibr1*, *ibr10* and *ibr1 ibr3 ibr10* mutants are fully insensitive to exogenous IBA in our growth conditions, thus making impossible to genetically address the putative dependency of IBR1 and IBR10 to glutathione levels. However, we noticed that in *ech2* and *ibr3* mutants, LR are still hyposensitive to IBA in the presence of BSO. Again, this suggests that ECH2 and IBR3 are not regulated in a glutathione-dependent manner. In order to detect an eventual defect in gene expression in glutathione-deficient plants, we analysed the expression level of genes involved in IBA-to-IAA conversion (Figure 6B). Surprisingly, *ECH2*, *IBR1*, *IBR3*, *IBR10*, *AIM1* and *PED1* genes were all moderately (10 to 50 % increase) but consistently upregulated in *cad2* mutant compared to the wild-type. In any case, this does not allow us to identify any target that could be transcriptionally down-regulated upon glutathione depletion. To complete this study and know if IBA conversion is affected, we tried to measure IBA-to-IAA conversion rate but unfortunately failed to obtain such data.

**Figure 6.**
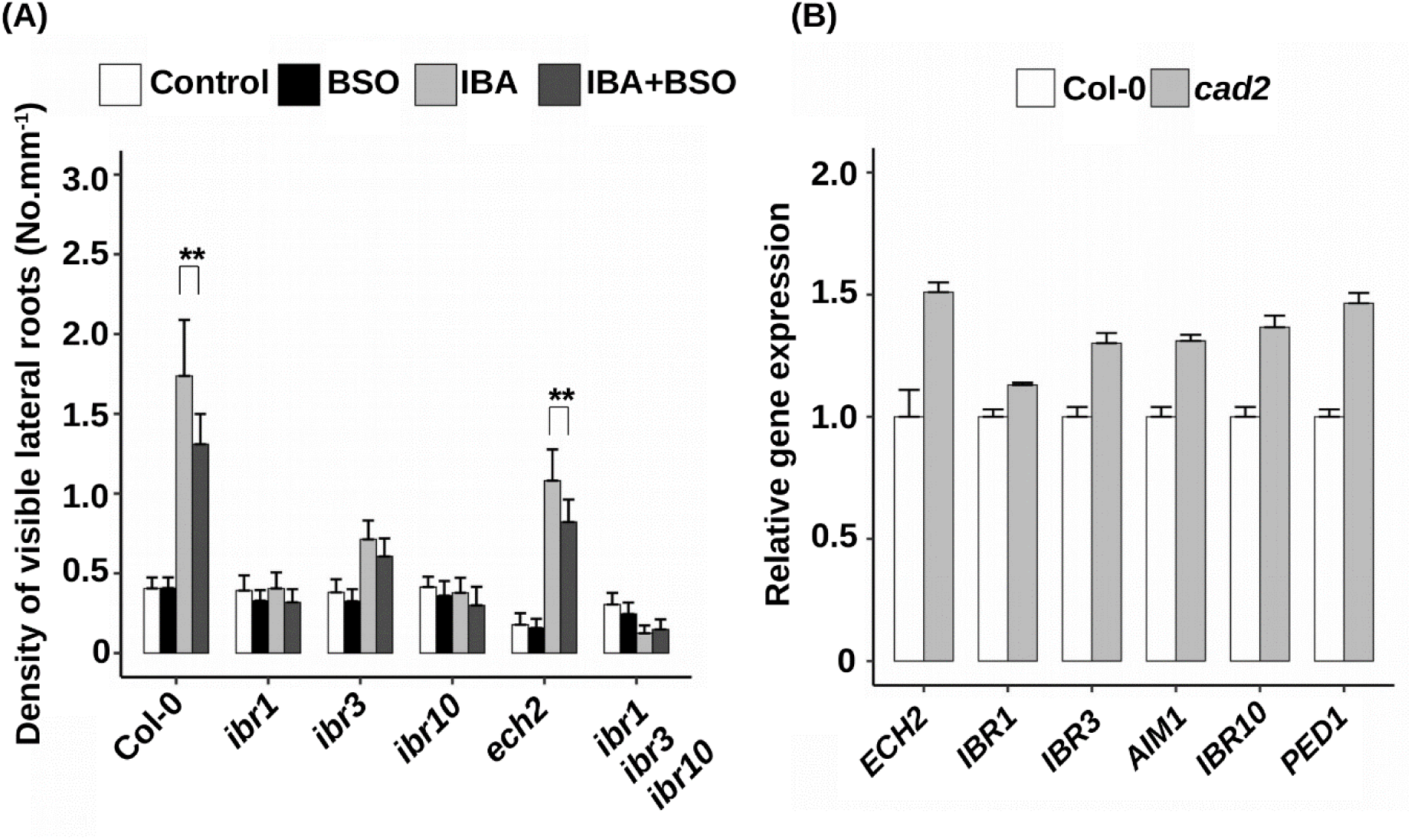
IBA conversion to IAA does not display sensitivity to glutathione. A. Emerged lateral root density of 10-day-old wild-type (Col-0), *ibr1*, *ibr3*, *ibr10*, *ibr1 ibr3 ibr10* or *ech2* plants grown on standard ½ MS medium, or in the presence of different combinations of BSO 0.5 mM and IBA 10 µM (n≥15). B. Expression levels (qRT-PCR) of *IBR1*, *IBR3*, *IBR10*, *ECH2*, *AIM1* and *PED1* genes in *cad2* relative to wild-type 8-day-old plants grown on standard ½ MS medium. Data represent three independent replicates with 20 plants per replicate. Histograms represent the mean and error bars represent the standard deviation. Asterisks indicate a significant difference, based on a two-tailed Student t-test (*P<0,01; **P<0,001).

To conclude, we carefully examined most of the known components of IBA homeostasis and response pathways, but none of them seem to be the target of glutathione-dependent regulation.

### IBA and glutathione control RH responses to phosphate deprivation

We have shown that LR and RH responses to exogenous IBA is dependent on glutathione levels. We then wanted to understand the physiological significance of such control. Since it had recently been shown that root-cap derived auxin regulates RH responses to phosphate deprivation, we decided to investigate RH responses to diverse environmental stimuli. Hence, we analysed RH elongation rate in response to phosphate starvation and to *Mesorhizobium loti* (*M. loti*), a well-described PGPR. We observed that both stimuli increase RH length in wild-type plants (Figure 7, A and B). Both *cad2* and *ibr1 ibr3 ibr10* mutants display wild-type like RH response to *M. loti*, suggesting that neither glutathione nor IBA participate in RH elongation response to PGPR (Figure 7A).

**Figure7.**
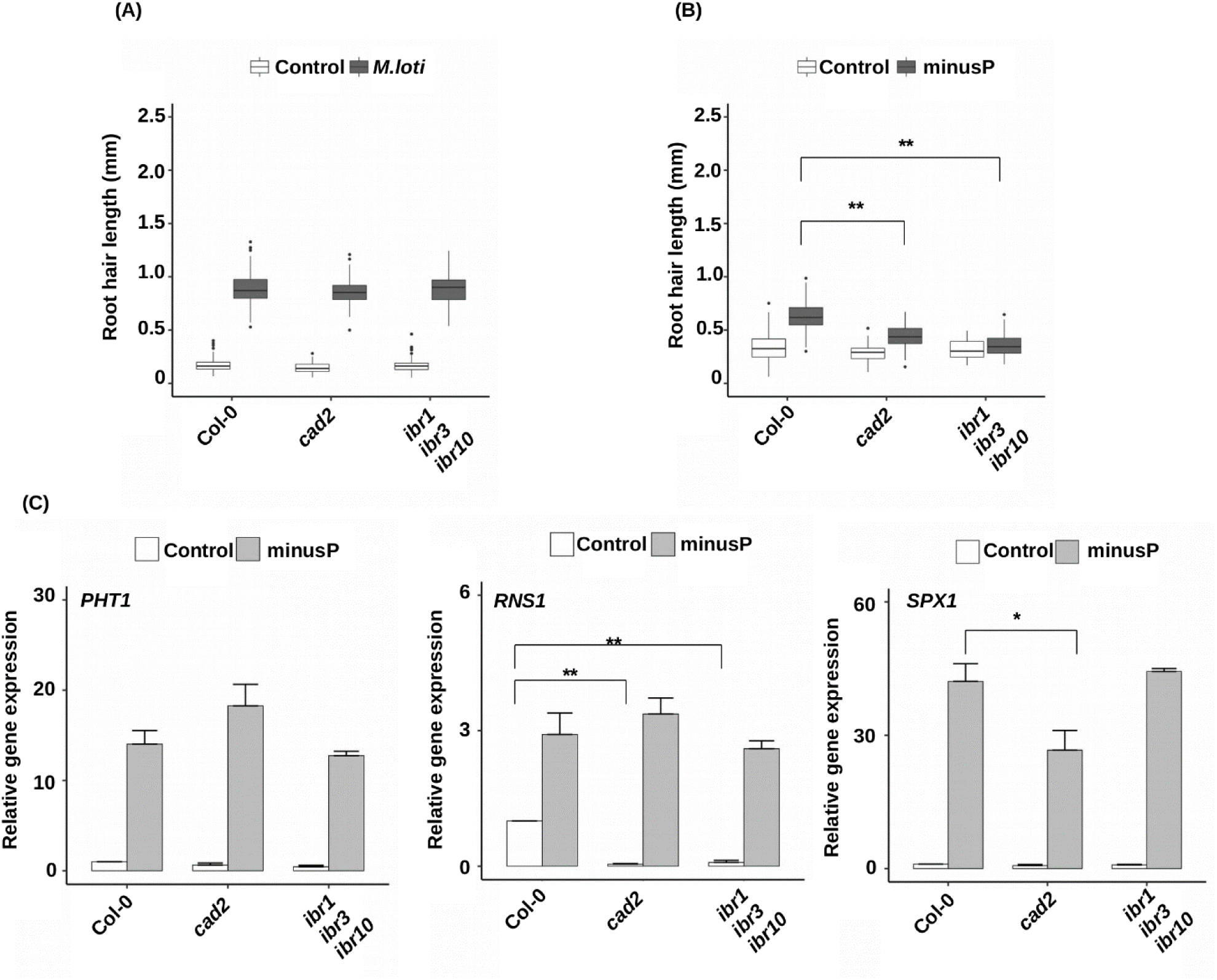
Glutathione and IBA are specifically required for RH growth response to low phosphate. A. RH length in 8-day-old wild-type (Col-0), cad2 or ibr1 ibr3 ibr10 plants grown on standard ½ MS medium in the absence or the presence of the PGPR *Mesorhizobium loti* (see material and methods). B. RH length in 8-day-old wild-type (Col-0), *cad2* or *ibr1 ibr3 ibr10* plants grown on 3/10 MS medium supplemented with 500 µM sodium phosphate (control) or 500 µM sodium chloride (minusP). C. Expression levels (qRT-PCR) of *PHT1;4*, *RNS1* and *SPX1* low-phosphate responsive genes, relatively to *ACT2* control gene, in wild-type (Col-0), *cad2* or *ibr1 ibr3 ibr10* 8-day-old plants grown on 1/3 MS medium supplemented with 500 µM NaH_2_PO_4_ (control) or 500 µM NaCl (minusP). Data represent three independent replicates with 20 plants per replicate and are normalized relatively to the wild-type value in control condition. Histograms represent the mean and error bars represent the standard deviation. Asterisks indicate a significant difference, based on a two-tailed Student t-test (*P<0,01; **P<0,001).

In contrast we observed that *cad2* mutant is clearly hyposensitive to phosphate deprivation, and that RH elongation response to phosphate starvation is almost fully abolished in *ibr1 ibr3 ibr10* mutant (Figure 7B). We also confirmed the decreased glutathione levels in *cad2* mutant while *ibr1 ibr3 ibr10* mutant does not alter glutathione content and redox status (Supplemental Figure 5). Interestingly, phosphate deprivation does not significantly affect total glutathione content in plants but increases the GSH/GSSG ratio.

We wondered if *cad2* and *ibr1 ibr3 ibr10* mutants affect the general response to phosphate deficiency or specifically the response of root hairs to this abiotic stress. We therefore quantified the expression of marker genes known to be induced in response to phosphate deprivation. Figure 7C shows that the induction of the expression of marker genes in response to phosphate starvation is not abolished in *cad2* and *ibr1 ibr3 ibr10* mutants. We only noticed a slight down-regulation of *SPX1* in *cad2* mutant. As *SPX1* encodes a negative regulator of phosphate starvation responses, this could explain why other responsive genes are slightly up-regulated in *cad2*.

We can conclude that both IBA conversion to IAA and glutathione are required for proper root hair response to phosphate starvation, without affecting general responses to this abiotic stress.

## DISCUSSION

### Glutathione regulation of IBA homeostasis or conversion to IAA in the root cap

In this work, we first report that root hair and lateral root responses to exogenous IBA, but not IAA, are impaired by glutathione deficiency. More precisely, the glutathione-dependent control of IBA response affects early steps of LR development, either founder cells specification or their activation to divide and form a LR primordium. We also show that low glutathione levels alter IBA-dependent auxin signalling in the basal meristem, where RH elongation occurs and LR founder cells are specified. Previous works have reported the central role of the root cap for IBA-to-IAA conversion in the context of LR founder cell specification, and the role of IAA transport from the root cap to the basal meristem for controlling RH elongation (Bhosale et al., 2018; De Rybel et al., 2012; Giri et al., 2018). Finally, we show in this manuscript that transporters known to be involved in the apico-basal flux of IAA to the basal meristem, namely AUX1 and PIN2, are not the targets of the glutathione-dependent control. All these data strongly suggest that glutathione modulates IBA homeostasis or IBA-to-IAA conversion in the root cap, although it does not exclude that the same regulation can also occur in additional tissues.

All the phenotypes reported here concern root responses to exogenous IBA and one can wonder if the glutathione-dependent regulation normally occurs in physiological conditions, modulating pathways relying on endogenous IBA. Because of our scarce knowledge of IBA metabolism and the absence of genetic tools concerning IBA biosynthesis, it is rather difficult to address such question. However, we accumulated several clues that, taken together, strongly support the importance of glutathione for the control of endogenous IBA pathways. First, we observe in *cad2* mutant seedlings grown under standard conditions (*i.e.* without exogenous IBA) a general increase in the expression of all genes involved in IBA-to-IAA conversion (Figure 6B). This might reveal a feedback mechanism that would report an excess of IBA or a depletion in IBA-derived IAA in low glutathione conditions. Secondly, we observe that both *ugt* and *pdr* double mutants are hypersensitive to exogenous IBA (Figures 5B and 5C), which is consistent with the already reported overaccumulation of endogenous free IBA in corresponding single mutants (Strader and Bartel, 2009; Ruzicka et al., 2010; Tognetti et al., 2010). Interestingly, it looks like BSO has more effect in both mutant genotypes than for the wild-type plants, reducing IBA response by 20 to 25% in wild type plants, but by 28 to 33% in *pdr* and *ugt* double mutants, respectively (Figures 5B and 5C). This supports a role for glutathione in modulating responses to endogenous IBA. Finally, we show that both *cad2* and *ibr1 ibr3 ibr10* triple mutant display hyposensitivity in their RH response to phosphate, but not to *Mesorhizobium loti*. In addition both are not impaired in the general plant response to phosphate deprivation, but only in the RH elongation response. This set of arguments proves that both endogenous IBA pathway and glutathione mediate RH response to phosphate starvation, and strongly suggests that the glutathione-dependent regulation of endogenous IBA pathway is modulating the RH response.

### Auxins fine-tuning of root architecture in response to phosphate availability

Although well-known to play important roles in plant development, the function of IBA as a source of IAA relative to other IAA sources, such as *de novo* synthesis or conjugated forms, is still very mysterious and we might wonder “Why plants need more than one type of auxins?” (Frick and Strader, 2018; Simon and Petrášek, 2011). A previous work has reported that IBA regulates both plant development and resistance to water stress, suggesting an important function for IBA in adapting plant development to abiotic stress (Tognetti et al., 2010). However, to our knowledge, our work is the first to clearly demonstrate that IBA-dependent pathway is necessary for IAA to adapt some developmental pathway, *i.e.* root hair elongation, to a specific abiotic stress (phosphate starvation).

In their recent work, Bhosale et al. (2018) already revealed the increase in IAA levels in the root cap in response to low external phosphate, necessary for RH elongation. However, because they observe a significant increase in *TAA1* (*TRYPTOPHAN AMINOTRANSFERASE OF ARABIDOPSIS 1*) gene expression in such conditions, they hypothesized that the increase in IAA levels in the root cap is due to the induction of Indole-3-Pyruvic Acid (IPyA)-dependent pathway, the main one responsible for *de novo* IAA synthesis in Arabidopsis (Mashiguchi et al., 2011). Intriguingly, RH elongation response to external low phosphate is almost completely abolished in both *ibr1 ibr3 ibr10* and *taa1* mutants (Figure 7B and Bhosale et al., 2018: Figures 2b and 2c). Could we imagine that two independent routes of IAA synthesis, *i.e.* the *TAA1*-dependent *de novo* synthesis and the IBA-to-IAA conversion, are both critical and induced in the root cap in response to phosphate deprivation? IBA can itself be derived from IAA and we can imagine that an increase in IBA-to-IAA conversion would require an increase in IBA levels, and therefore more IAA to supply the IBA stock. However, thinking to a root cap increase in IAA to increase IBA in order to increase IAA gets almost no sense. Thus, we suggest that external low phosphate induces *TAA1* expression in the whole plant to increase global available auxin levels on the one hand, and in the other hand activates the local IBA-to-IAA conversion in the root cap to ensure a local appropriate developmental response.

However, we must remind that we are working *in vitro* with homogenous media containing a given concentration of phosphate. A very elegant recent work has reported that, when subjected to an heterogenous environment, using a Dual-flow-root chip, a single root increases RH length in the medium having the highest phosphate concentration (Stanley et al., 2018). This observation contrasts with the induction of RH growth by low phosphate in homogenous *in vitro* media. RH adaptation to phosphate probably results from the crosstalk between a systemic information based on overall available phosphorus in the plant, and a local information based on phosphate availability in the environment, that we cannot differentiate in our *in vitro* homogenous conditions. Such dual control has been extensively documented for root responses to another important nutrient, nitrogen (Ruffel and Gojon, 2017). We might therefore envisage that both sources of IAA are differentially regulated by systemic and local signals. This might ensure a very acute dosage of free IAA to optimize root adaptation to the plant metabolism and environment. In such context, glutathione could be a central regulator of local IBA-to-IAA adaptation to external phosphate in the root cap.

### Unravelling the glutathione-dependent regulation of IBA pathway

Although we explored most of known components of IBA pathways, we were not able to identify how glutathione regulates IBA homeostasis or conversion to IAA. This can first be explained by the difficulties to work with IBA which is present in low amounts compared to IAA in plants, and is therefore difficult to detect and quantify (Frick and Strader, 2018). Moreover, many components of the IBA pathways remain to be identified. For sure, the development of new tools, such as IBA probes or mutants in IBA biosynthesis would be very helpful to depict the important roles played by this auxin in plants. The other problem we faced is the complex multifaceted roles of glutathione in cells, that prevented us to try to decipher the mechanism from the glutathione starting point. The only clue we get is that IBA hyposensitivity is specific to plants with reduced amounts of glutathione but does not occur in plants having more general redox problems (Figure 3). This suggests that IBA regulation by glutathione may act through glutathionylation or via the activity of thiols reductases that specifically depend on glutathione, such as glutaredoxins. One hypothesis would be that IBA, or a precursor, could itself be glutathionylated, which would be responsible for its storage or transport. In Arabidopsis, the Tau class of Glutathione-S-Transferases (GSTU) have been shown to glutathionylate some fatty acids, ranging from short to long acyl chains (Dixon and Edwards, 2009). Similarly, GSTU19 and GSTF8 have been proposed to glutathionylate 12-oxo-phytodienoic acid (OPDA), hence allowing its translocation from the chloroplast to the peroxisome where it is converted to Jasmonic Acid (Davoine, 2006; Dixon and Edwards, 2009).

Because IBA-to-IAA conversion occurs in the peroxisome, we might also envisage that the glutathione-dependent regulation is not specific to IBA pathways but affects the peroxisomal machinery. One of the main peroxisomal functions is the β-oxidation of fatty acids, which is essential to supply energy during seed germination. We never observed any problem for seed germination in *cad2* or *pad2* mutants, in contrast to mutants in peroxisomal functions which generally strongly affect seed germination. However, we cannot exclude that the glutathione-dependent regulation only occurs in the root cap and therefore mainly affects IBA-ot-IAA conversion although regulating the peroxisome machinery. Among the PEROXINS proteins, many participate in the peroxisomal matrix protein import machinery (Cross et al., 2016). PEX5 is a central cargo protein that recognizes proteins targeted with the specific PTS1 (Peroxisomal Targeting Signal 1) signal peptide and import them into the peroxisome. PEX7 recognizes proteins harbouring another signal peptide (PTS2), and then binds to PEX5 for import into the peroxisome. Interestingly, previous works on human and *Pichia pastoris* PEX5 revealed that its activity and oligomerization depend on a redox switch affecting a conserved N-terminal Cystein (Ma et al., 2013; Apanasets et al., 2014). However, such regulation is not specific to glutathione but rather depends on the redox status and the content in ROS of peroxisomes.

Finally, the last hypothesis would be that glutathione regulates IBA homeostasis or conversion to IAA via a yet unidentified or not assayed component. New attempts in quantifying IBA-to-IAA conversion would be helpful in order to confirm this hypothesis. Concerning the IBA-to-IAA β-oxidation pathway in the peroxisome, PED1 has been reported to harbour a redox-sensitive switch affecting a conserved cysteine and that participates in activating the enzyme when reduced (Pye et al., 2010). In contrast to most of the other enzymes involved in the conversion pathway, PED1 is targeted to the peroxisome through a PTS2 signal peptide. It is interesting to notice that *pex5-1*, in contrast to *pex5-10*, is not altered in PTS1-dependent import but only in PTS2-dependent import of proteins into the peroxisome (Woodward and Bartel, 2004; Khan and Zolman, 2010). *pex5-1*, although not displaying any phenotype during germination, is highly affected in IBA responses. This might reveal that IBA to IAA conversion is highly dependent on a PTS2-targeted protein, and PED1 could therefore be a good candidate to assay. However, PED1 is not specific to IBA-to-IAA pathway since it acts in almost all β-oxidation pathways occurring in the peroxisome. We could imagine a local glutathione-dependent regulation of PED1 activity specifically in the root cap.

In conclusion, our work points out the important role of glutathione in both controlling IBA-to-IAA conversion required for proper root development and mediating developmental responses to nutrient deprivation. This opens novel perspectives in understanding how redox homeostasis can signal and integrate environmental constraints to trigger proper developmental adaptations via the regulation of morphogenetic hormonal pathways.

## MATERIAL AND METHODS

### Plant Material

Wild-type and mutants plants used throughout this study are in the *Arabidopsis thaliana* Col-0 ecotype. The following mutants were all previously published and available in our teams : *cad2-1* (Howden et al., 1995), *pad2-1* (Parisy et al., 2007), *zir1* (Shanmugam et al., 2012), *gr1* (Mhamdi et al., 2010), *ntra ntrb* (Bashandy et al., 2010), *cat2* (Bueso et al., 2007), *ibr1-2* (Zolman et al., 2008), *ibr3-1* (Zolman et al., 2007), *ibr10-1* (Zolman et al., 2008), *ibr1 ibr3 ibr10* (Zolman et al., 2008), *ech2* (Strader et al., 2011), *aux1-21* (Bennett et al., 1996), *pin2/eir1-1* (Roman et al., 1995). *pdr8/pen3-4* (Stein et al., 2006), *pdr9-2* (Ito and Gray, 2006), *ugt74e2* (Tognetti et al., 2010), and *ugt74d1* (Jin et al., 2013) are previously described Salk T-DNA insertion lines (Salk_000578, Salk_050885, Salk_091130 and Salk_011286, respectively) ordered from the Nottingham Arabidopsis Stock Centre (NASC; http://arabidopsis.info/). The double mutants *pdr8 pdr9* and *ugt74d1 ugt74e2* were obtained by crossing the respective single mutants, and by selecting the double homozygous among F2 plants.

### Plant Cultures

Seeds were surface sterilized by constant agitation with 0.05% SDS in 70% (v/v) ethanol by 20 minutes, then washed three times with ethanol 95% (v/v) and dried on sterile paper. Seeds were placed on square plates containing 50 mL of ½ Murashige and Skoog medium (MS) with 0.5 g.L^-1^ MES and 0.8 % (w/v) plant agar (Duchefa Biochemie) without sucrose unless indicated. For the experiments regarding phosphate deprivation, we used three-tenth-strength (3/10) MS supplemented with 500 µM NaCl (phosphate deprivation) or 500 µM NaH2PO4 (control conditions) as previously published (Arnaud et al., 2014). For *Mesorhizobium loti* experiments, seeds were grown on standard ½ MS medium for 5 days, then transferred to new plates containing 0.1 OD of inoculum (Poitout et al., 2017) for 4 extra days before phenotyping. All plates were incubated vertically at 20°C with 160 µE.m^-2^.s^-1^ light intensity and a 16h-light/8h-dark regime.

### Phenotypic Analyses of the Root System

Lateral root density was quantified on 10-day-old seedlings; this consisted in counting the number of visible lateral roots emerged and dividing by the length of the primary root section between the first and the last visible lateral roots. Lateral root primordia density was calculated in the same way on 6-day-old seedlings using a light microscope (Axioscop2, Zeiss) for counting. For gravistimulation experiments, LRP initiation was forced by changing the orientation of the vertical plates by 90°. LRP stages were observed 48h later, using a light microscope (Axioscop2, Zeiss). For root hairs lengths quantification, plates were photographed using a camera (DFC425C, Leica) sited on a stereomicroscope (MZ12, Leica). Primary root elongation rate was quantified between day 8 and day 10. Lengths were quantified from pictures using the public domain image analysis program ImageJ 1.52i (https://imagej.nih.gov/ij/) and its NeuronJ plugin (Meijering et al., 2004).

### Glutathione Measurements

Total glutathione content of 8-day-old seedlings was determined using the recycling enzymatic assay (Rahman et al., 2007). The method consists in the reduction of GSSG by glutathione reductase enzyme and NADPH to GSH. GSH levels are determined by the oxidation of GSH by 5,5′-dithio-bis(2-nitrobenzoic acid) (DTNB), that produces a yellow compound 5′-thio-2-nitrobenzoic acid (TNB), measurable at 412 nm (Rahman et al., 2007). Briefly, 100 mg of fresh plant material ground in liquid nitrogen was extracted in 0.5 mL 0.1 M Na phosphate buffer, pH 7.6, and 5 mM EDTA. After micro-centrifugation (10 min, 9000g), total glutathione in 0.1 mL of the supernatant was measured by spectrophotometry in a 1 mL mixture containing 6 mM dithiobis(2-nitrobenzoic acid) (Sigma-Aldrich), 3 mM NADPH, and 2 units of glutathione reductase from Saccharomyces cerevisiae (Sigma-Aldrich). Glutathione-dependent reduction of dithiobis(2-nitrobenzoic acid) was followed at 412 nm. Total glutathione levels were calculated using the equation of the linear regression obtained from a standard GSH curve. Oxidized glutathione (GSSG) was determined in the same extracts after derivatization of reduced GSH. Derivatization of 100 mL plant extract was performed in 0.5 mL 0.5 M K phosphate buffer, pH 7.6, in the presence of 4 mL 2-vinylpyridine (Sigma-Aldrich) during 1 h at room temperature. After extraction of the GSH-conjugated 2-vinyl pyridine with 1 volume of diethylether, the GSSG was measured by spectrophotometry as described for total glutathione.

### Confocal Analyses

Auxin response analyses using *DR5:n3EGFP/DR5v2:ntdTomato* (Liao et al., 2015) and the study of the redox state of glutathione with roGFP2 line (Meyer et al., 2007) were performed by using a confocal laser scanning microscope LSM 700 (Zeiss). Images were acquired in 16-bits using a 10x EC Plan Neofluar objective (Zeiss). Settings were based on the respective methods previously published (Liao et al., 2015; Meyer et al., 2007) with minor modifications. Excitation of roGFP2 was performed at 488 and 405 nm and a bandpass (BP 490-555 nm) emission filter was used to collect roGFP2 signal. For background subtraction, signal was recovered using a BP 420-480 nm emission filter during excitation at 405 nm.

For *DR5:n3EGFP/DR5v2:ntdTomato* analysis, seeds were grown vertically on standard ½ MS medium supplemented or not with BSO 0.5mM (pre-treatment). 7-day-old seedlings were transferred for 24 hours to plates containing the appropriate treatments (IBA 10µM, IAA 50nM), still combined or not with BSO 0.5mM, according to the pre-treatment. Regarding the analyses with roGFP2, seeds were grown vertically for 8 days in ½ MS medium containing or not IBA 10µM.

Pictures analyses and quantifications were performed as previously described for both probes (Meyer et al., 2007; Liao et al., 2015), using the public domain image analysis program ImageJ 1.52i (https://imagej.nih.gov/ij/).

### Histochemical Localization of GUS Activity

Plants were fixed in 80% (v/v) acetone at 20 °C for 20 minutes, then washed with buffer solution, containing 25 mM Na_2_HPO_4_, 25mM NaH_2_PO_4_, 2% Triton X-100 (v/v), 1 mM K_3_Fe(CN)_6_, and 1 mM K_4_Fe(CN)_6_. Thereafter, staining was performed at 37 °C in the buffer solution containing 2mM of X-Gluc (5-bromo-4-chloro-3-indolyl-β-D-glucuronidase) as substrate (Jefferson et al., 1987), after 1 min vacuum infiltration. The reaction was stopped by changing the seedlings to ethanol 70% (v/v). Pictures were collected using a light microscope (Axioscop2, Zeiss).

### Gene Expression Quantification

Total RNA was extracted using TriZol reagent (GE Healthcare, UK), and the RNA was purified with RNeasy Plant Mini Kit (Promega, USA), according to the manufacturer protocols. cDNA was subsequently synthesized using the GoScript™ Reverse Transcription System (Promega, USA). Quantitative real-time PCR was done using Takyon™ No Rox SYBR® MasterMix blue dTTP (Eurogentec, Belgium) and the LightCycler 480 (Roche, Switzerland). Primers used are presented in Supplemental Table 1. All reported results are presented normalized with the *ACTIN2* control gene but behave similarly if normalized with *ACTIN7* or *GAPDH* control genes.

### Auxin Accumulation Assays

5-mm root tips from 8-day-old seedlings were excised and incubated in 40 µL uptake buffer (20 mM MES, 10 mM sucrose, and 0.5 mM CaSO_4_, pH 5.6) for 15 min at room temperature. Root tips were then incubated for one hour in uptake buffer containing 25 nM [^3^H]-IAA (20 Ci/mmol; American Radiolabeled Chemicals) or [^3^H]-IBA (25 Ci/mmol; American Radiolabeled Chemicals) prior to washing with three changes of uptake buffer. Root tips were then placed in 3 mL CytoScint-ES liquid scintillation cocktail (MP Biomedicals) and analyzed by scintillation counting.

### Data Replicability and Statistical Analyses

All the experiments presented in this study illustrate results obtained in at least three independent biological repetitions. For *in vitro* phenotyping experiments, each biological repetition consisted in two technical replicate plates per condition, each containing around 12 seeds for the mutant and 12 for the appropriate control, sown side by side. Statistical analyses were performed as indicated in Figure legends.

## Accession Numbers

Sequences from this article can be found in the Arabidopsis Genome Initiative database under the following accession numbers : *GSH1* (*At4g23100*), *GR1* (*At3g24170*), *NTRA* (*At2g17420*), *NTRB* (*At4g35460*), *CAT2* (*At4g35090*), *AUX1* (*At2g38120*), *PIN2* (*At5g57090*), *PDR8/PEN3* (*At1g59870*), *PDR9* (*At3g53480*), *UGT74D1* (*At2g31750*), *UGT74E2* (*At1g05680*), *IBR1* (*At4g05530*), *IBR3* (*At3g06810*), *IBR10* (*At4g14430*), *ECH2* (*At1g76150*), *AIM1* (*At4g29010*), *PED1* (*At2g33150*), *PHT1;4* (*At2g38940*), *RNS1* (*At2g02990*), *SPX1* (*At5g20150*), *ACT2* (*At3g18780*).

## Supplemental Data

The following materials are available in the online version of this article.

**Supplemental Figure 1.** Glutathione does not affect primary root response to IBA.

**Supplemental Figure 2.** Glutathione regulates LR responses to IBA.

**Supplemental Figure 3.** IBA does not alter redox status of GSH in root tips.

**Supplemental Figure 4.** RH response to IBA in mutants affected in IBA homeostasis.

**Supplemental Figure 5.** Glutathione content in response to low phosphate.

**Supplemental Table 1.** List of primers used for qPCR analyses.

## ACKNOWLEDGMENTS

We want to thank Edouard Jobet for his technical support for qPCR experiments. We thank the “Bio-Environnement” technological UPVD platform (Région Occitanie, CPER 2007-2013 Technoviv, CPER 2015-2020 Technoviv2), for access to the confocal microscope and qPCR facilities. This work was supported by the “Centre National de la Recherche Scientifique” and the “Université de Perpignan Via Domitia”. This study is set within the framework of the “Laboratoires d’Excellence (LabEx)” TULIP (ANR-10-LABX-41). JA T-H is supported by a PhD grant from “Consejo Nacional de Ciencia y Tecnología” (CONACYT:254639/411761).

## AUTHOR CONTRIBUTIONS

J.A. T.-H., and C. B. designed the experiments and analysed data. J.A. T.-H. performed most of the experiments. L.B. assisted with crosses and generation of double mutants; C.B. and J.A. T.-H. wrote the manuscript; J.-P. R. and L.C. S. consulted on the study and revised the manuscript. J.-P. R. and C.B. supervised the project.

---

**Supplemental Figure 1.** Glutathione does not affect primary root response to IBA.

This supplemental Figure supports Figure 1.

A. Pictures of representative 8-day-old wild-type, *cad2*, *pad2*, and *zir1* plants grown on standard ½ MS medium supplemented or not with 10 µM IBA or 50 nM IAA. Wild-type plants grown on 0.5 mM BSO are also presented.

B. Quantification of primary root elongation rate of 8-day-old wild-type (Col-0), *cad2* or *pad2* mutant plants grown on standard ½ MS medium (Control), or in the presence of 50 nM IAA (IAA) or 10 µM IBA (IBA). Wild-type plants in the presence of 0.5 mM BSO (BSO) were also assayed (n≥12).

Histograms represent the mean and error bars represent the standard deviation. Asterisks indicate a significant difference, based on a two-tailed Student t-test (*P<0,01; **P<0,001).

**Supplemental Figure 2.** Glutathione regulates LR responses to IBA.

This supplemental Figure supports Figure 1.

A. Emerged LR density of 10-day-old wild-type plants grown on standard ½ MS medium (control), or in the presence of different concentrations of BSO, in the absence or the presence of IBA 10 µM (n≥15).

B. Emerged LR density of 10-day-old wild-type and *cad2* plants grown on standard ½ MS medium (control), or in the presence of IBA 10 µM and 5 mM GSH (n≥15).

Histograms represent the mean and error bars represent the standard deviation. Asterisks indicate a significant difference, based on a two-tailed Student t-test (*P<0,01; **P<0,001).

**Supplemental Figure 3.** IBA does not alter redox status of GSH in root tips.

This supplemental Figure supports Figure 3.

A. Kinetics of roGFP2 reporter response to 1 mM H_2_O_2_ then to an additional 10 min treatment with 10 mM DTT. The ratio between the oxidised roGFP2 signal (excitation at 405 nm) and the reduced roGFP2 signal (excitation at 488 nm) is used to monitor the glutathione redox status.

B. Quantification of roGFP2 ratio in root tips from wild-type plants grown on standard ½ MS medium or in the presence of 10 µM IBA for 8 days.

Data represent the mean and error bars represent the standard deviation (n≥9).

**Supplemental Figure 4.** RH response to IBA in mutants affected in IBA homeostasis.

This supplemental Figure supports Figure 5.

RH length of 8-day-old wild-type (Col-0), *ugt74e2 ugt75d1* and *pdr8 pdr9* plants grown on standard ½ MS medium, or in the presence of different combinations of BSO 0.5 mM and IBA 10 µM (n≥50). Asterisks indicate a significant difference, based on a two-tailed Student t-test (**P<0,001).

**Supplemental Figure 5.** Glutathione content in response to low phosphate.

This supplemental Figure supports Figure 7.

Glutathione content in 8-day-old wild-type (Col-0), *cad2*, or *ibr1 ibr3 ibr10* seedlings grown on 1/3 MS medium supplemented with 500 µM NaH_2_PO_4_ (control) or 500 µM NaCl (minusP). Data represent the means of 3 biological repetitions. Reduced glutathione is represented in white (GSH) and oxidised glutathione in black (GSSG). Histograms represent the mean and error bars represent the standard deviation.

**Supplemental Table 1.** List of primers used for qPCR analyses.

